# Geometric-relationship specific transfer in visual perceptual learning

**DOI:** 10.1101/2023.12.07.570648

**Authors:** Qingleng Tan, Yuka Sasaki, Takeo Watanabe

## Abstract

Visual perceptual learning (VPL) is defined as long-term improvement on a visual task as a result of visual experience. In many cases, the improvement is highly specific to the location where the target is presented, which refers to location specificity. In the current study, we investigated the effect of a geometrical relationship between the trained location and an untrained location on transfer of VPL. We found that significant transfer occurs either diagonally or along a line passing the fixation point. This indicates that whether location specificity or location transfer occurs at least partially depends on the geometrical relationship between trained location and an untrained location.

## Introduction

### Visual Perceptual Learning

(VPL) refers to a long-term enhancement in visual task performance as a result of visual experience (Chang et al., 2014; Gold & Watanabe; Sasaki et al., 2010; Sasaki & Watanabe, 2012; Watanabe & Sasaki, 2015; Frank et al, 2020; Frank et al, 2022; Yotsumoto & Watanabe, 2008; Sagi & Tanne, 1984; Dosher & Lu, 1998; Fahle & Poggio, 2002; Seitz et al., 2005; DeLoss et al., 2015; Roelfsema et al, 2010; Yamada et al, 2023). It has been found that VPL can improve our visual abilities to process a variety of features, including orientation (Bang et al., 2018; Schoups et al., 1995; Shibata et al., 2011; Shibata et al., 2012; Shibata et al., 2017; Tsushima & Watanabe, 2009), motion direction (Ball and Sekuler, 1987; Watanabe et al., 2001; Watanabe et al., 2002; Seitz and Watanabe, 2003), and texture (Karni and Sagi, 1991). VPL has been extensively studied particularly in the past two decades mainly because of its close links to cortical plasticity (Gilbert et al., 2001; Yang & Maunsell, 2004; Schoups et al., 2001; Law and Gold, 2008; Yotsumoto et al, 2008; Byers & Serences, 2014; Shibata et al, 2017). VPL is also regarded as a promising tool with which to improve degraded or declined perceptual abilities due to visual diseases (Levi & Polat, 1996; Levi, 2009) or aging (Andersen et al., 2010; Yotsumoto et al., 2014, Lemon & DeLoss, 2016). Thus VPL is regarded as a vital subject in visual science.

One of the most distinguishing characteristics of many types of VPL is its high specificity of the trained retinal location, which refers to **location specificity**. For instance, VPL of orientation or motion direction is largely confined to the location where the stimulus is presented (Karni & Sagi, 1992; Ball & Sekuler, 1987; Watanabe et al, 2002; Yotsumoto et al., 2008). Location specificity of VPL has been regarded as a manifestation of involvement of early visual areas, since neurons in these areas have relatively small receptive fields (Schoups et al., 2001).

However, there has been emerging evidence that location specificity of VPL is abolished by certain experimental procedures. Double training is an exemplar of those procedures. It has been found that location-specific contrast discrimination learning was rendered completely transferrable to the new location in which a subsequent orientation discrimination task was trained (Xiao et al., 2008; also see Liang et al. 2015 a & b), although whether transfer occurred or not depended on the training procedure (Hung & Seitz, 2014). Another study has found that training with an oriented target embedded in the texture, which usually results in location specificity of learning (Karni and Sagi, 1992), caused no location specificity if the target was followed by flanks in the same orientation of the target during training (Harris et al., 2012). These findings indicate that the location specificity of VPL of a task, which is shown as a result of usual training, can be abolished if a certain operation is added to training. This raises the possibility that standard training leads to location specific VPL but that some specific operations enables the location transfer.

Interestingly, whether VPL is location specific or not, these studies stand on the assumption that if VPL is not location specific, it should not occur in early stages. The results of the present study suggest that this assumption is not always the case. We found that whether location specificity or location transfer occurs depends on the geometric relationship between the trained location and an untrained location. This suggests that location transfer does not necessarily indicate high-level involvement in VPL.

### Experiment 1

We conducted orientation discrimination training and examined whether a geometric relationship between the trained and an untrained locations influences the degree of transfer of VPL.

## Methods

### Subjects

Twelve healthy subjects (aged 18 to 28) with normal or corrected-to-normal vision participated in the study. Subjects were naïve to the purpose of the study. Informed consent was obtained from all subjects.

### Stimuli and apparatus

Experiments were conducted in a dimly lit room. The stimuli were generated by matlab 2012b. The stimuli were presented on an 19-inch Pure Flat Sony Trinitron CRT, with a spatial resolution of 1024×768 and a refresh rate of 100 Hz. Subjects viewed the stimuli from a distance of 57 cm. Their head position was stabilized using a chinrest.

Visual stimuli were Gaussian windowed sinusoidal gratings (Gabors) with randomized phases (diameter: 5°; contrast: 0.9; spatial frequency: 2 cycles/°; orientation: around 15° right tilted from vertical (Fig. 1). They were located at 5° retinal eccentricity in one of the four visual quadrants.

**Figure 1.**
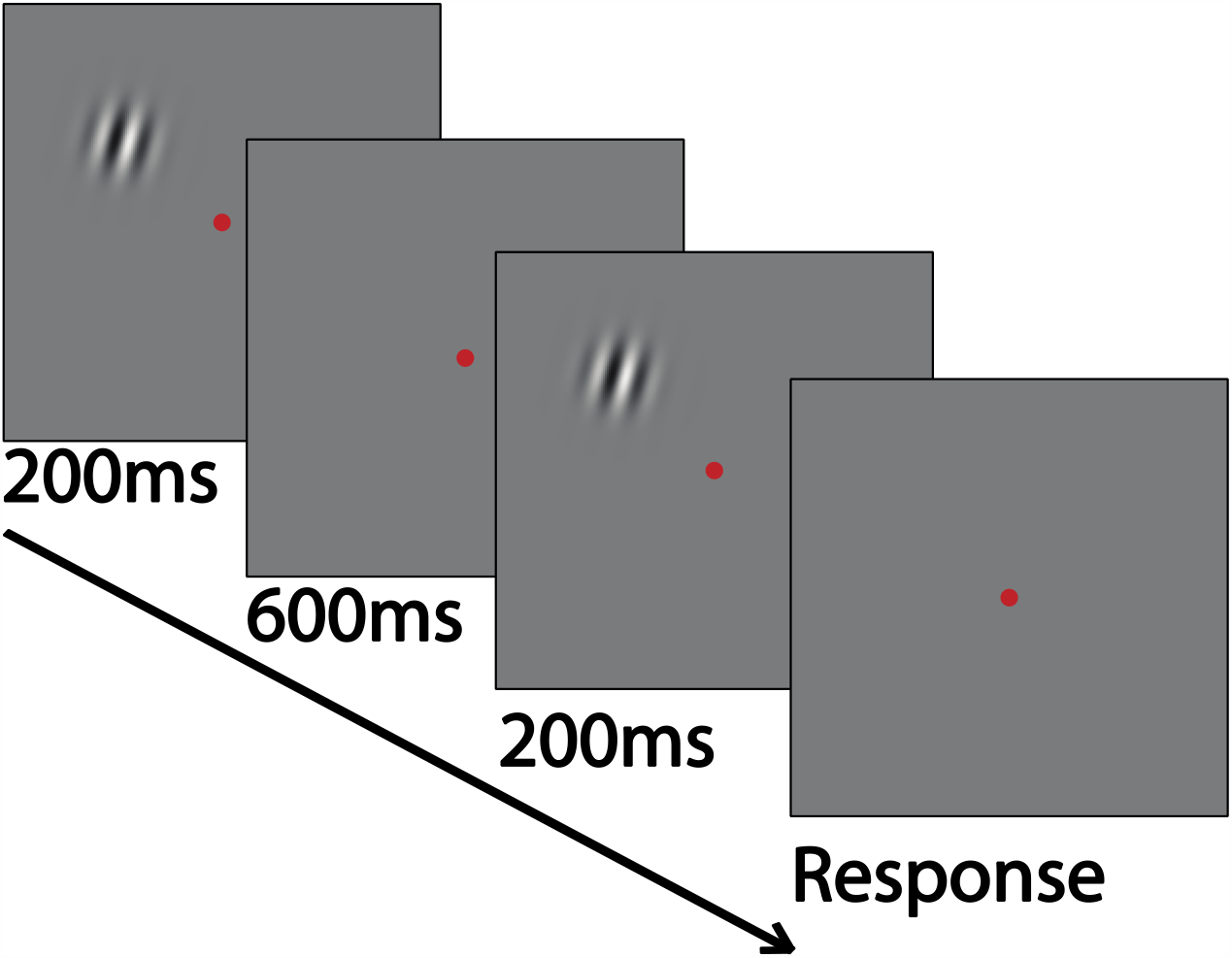
Procedure of a discrimination task.

### Procedures

The experiment consisted of the pre-test (day 1), training (day 2-7) and post-test (day 8). During the training phase, each subject underwent six daily training sessions on an orientation discrimination task. One daily session (about 1 h) consisted of 20 QUEST staircases of 40 trials (Watson and Pelli, <u>1983</u>). In each trial, test and reference stimuli were each presented for 200 ms and were separated by a 600 ms blank interval (Fig. 1). The temporal order of the presentations of test and reference stimuli was randomized across trials. Subjects were asked to make a forced choice judgment as to whether the second stimulus was rotated clockwise or counterclockwise from the first stimulus. A high-pitched tone was provided after a wrong response, whereas no tone was presented if the response was correct. The next trial began one second after subject’s response. The step size of the staircase varied trial by trial and was controlled by the QUEST staircase to estimate subject’s discrimination threshold (75% correct). Stimuli in the orientation discrimination task were presented consistently at the upper left quadrant throughout 6 days of training. During the test sessions, orientation discrimination thresholds were measured at the three locations (trained location, diagonal location and horizontal location) in a counter-balanced order. The diagonal location was defined as the point-symmetry location from the trained location with respect to the central fixation point. The horizontal location was defined as the line-symmetry location to the trained location with respect to the vertical line crossing the central fixation point. To measure an orientation discrimination threshold, 8 QUEST staircases (same as above) were completed for each location. Thresholds at each location were averaged within subjects. Subject’s performance improvement after training was calculated as (pre-test threshold-post-test threshold)/pre-test threshold ×100%.

## Results

Fig. 2 shows the mean threshold as a function of the stimulus location in the tests. We applied a 2-way ANOVA with locations (trained location, diagonal location vs. horizontal location) and test stages (pre-vs. post-tests) as main factors with repeated measures to thresholds. The results showed a significant main effect of test stages (F_(1,22)_= 15.68, *p*=0.02), while the main effect of locations was not significant (F_(2,22)_= 2.98, *p*=0.07). We also found a significant interaction between locations and test stages (F_(2,22)_= 8.47, *p*=0.02). We calculated percent improvements on three locations to reveal how training effects on performances at different locations. Paired t-test showed that, the improvements at the trained location (29.32%) and the diagonal location (22.00%) were significantly larger than the parallel location (9.74%)(t_11_ = 4.59, *p*<0.01, and t_11_ = 4.21, *p*<0.01, respectively, corrected for multiple comparisons). These results indicate that VPL of orientation discrimination significantly transferred to the diagonal location, whereas the degree of transfer to the horizontal location was not significant.

**Figure 2.**
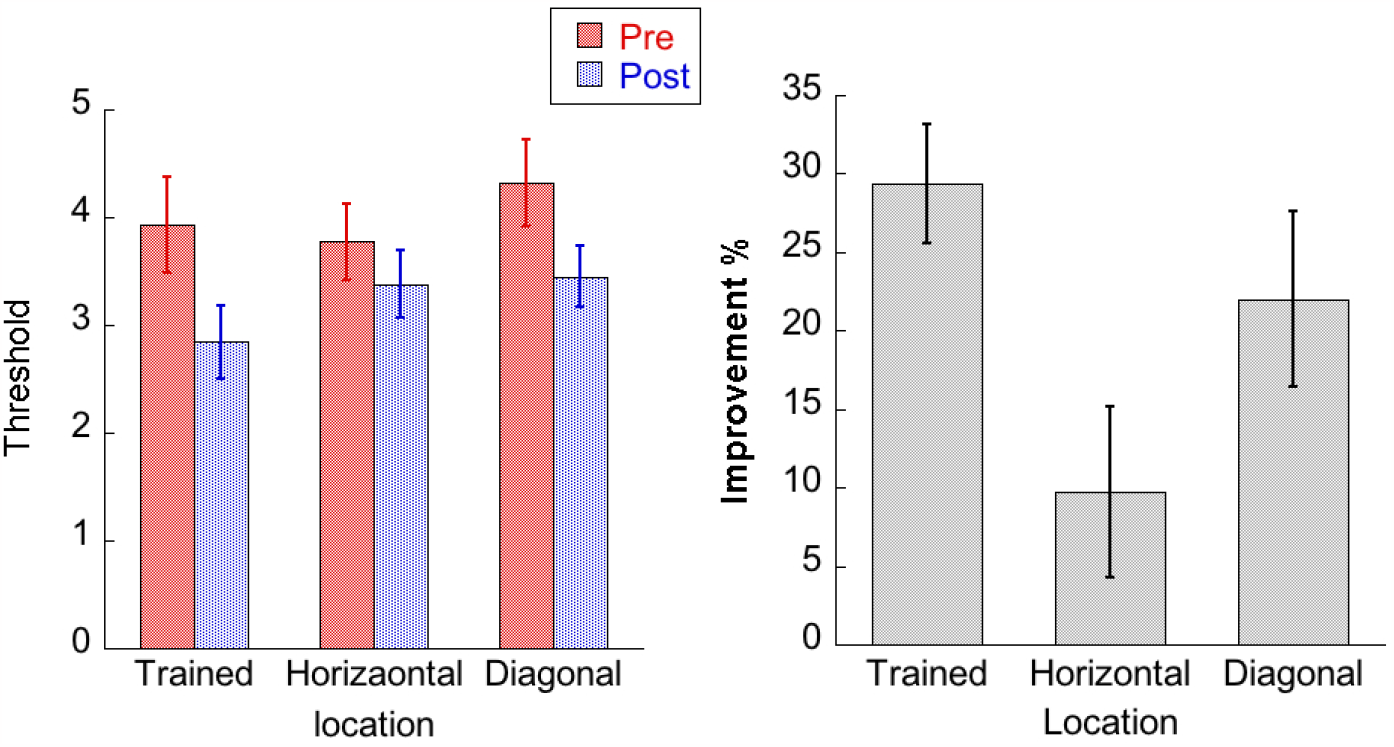
The result of Experiment 1

### Experiment 2

The result of Experiment 1 indicates that transfer of VL occurs to a diagonal location, but not to a horizontal location. To be noted, the distance between a trained location and a horizontal location is shorter than that between a trained location and a diagonal location. This indicates that whether transfer occurs or not depends on the geometrical relationship between the trained location and an untrained location. In Experiment 1, the trained grating and a stimulus in the diagonal location were symmetrically laid out with respect to the fixation point. However, the orientation of the grating in horizontal location is not. It has been found that symmetrical located gratings mutually enhance each other (Sasaki et al., 2005). A similar principle might have governed transfer, despite the fact that diagonal locations were more distant than the horizontal locations. To further investigate what geometrical relationship allows transfer to occur in Experiment 2, the orientation of grating in neither the trained nor the untrained location was arranged to be symmetrical.

## Methods

### Subjects

Eight new healthy subjects (aged 18 to 28) with normal or corrected-to-normal vision participated in the study. Subjects were naïve to the purpose of the study. Informed consent was obtained from all subjects.

### Stimuli

As shown in Fig. 3, the trained stimulus was presented 5° to the left to the fixation point. The diagonal location in a test stimulus was presented 5° below the fixation point. The horizontal location was presented 5° to the right to the fixation point. Thus, the line between the trained and the diagonal location did not pass the central fixation point, whereas the line between the trained and the horizontal location passed the fixation point.

**Figure 3.**
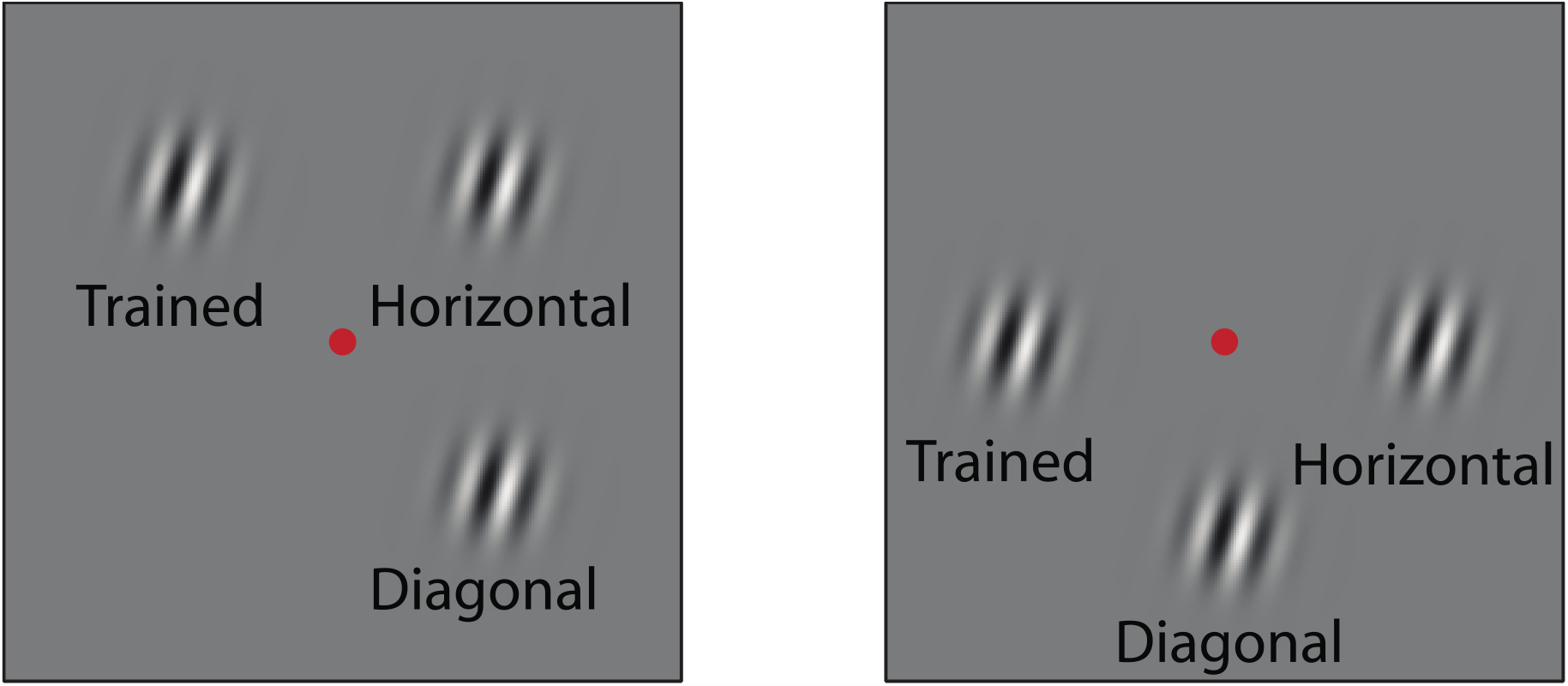
Schematic illustration of the trained location (T), diagonal location (D) and horizontal location (H) in test stages of Experiment 1 (left) and Experiment 2 (right).

## Results

The results are shown in Fig. 4. A two-way ANOVA was conducted in the same way as in Experiment 1. A significant main effect of test was found (F_(1,14)_= 59.41, *p*<0.01). A main effect of locations was also obtained (F_(1,14)_= 3.81, *p*=0.048). No significant interaction between location and test stage was found (F_(2,14)_= 1.94, *p*=1.18). These results indicate that transfer of VPL of orientation discrimination occurred from the trained to the horizontal location as well as from the trained to the diagonal location. We confirmed the result with further analysis on improvement at three locations. There was no significant difference among the improvements at three locations (F_(2,14)_=0.82, *p*=0.46).

**Figure 4.**
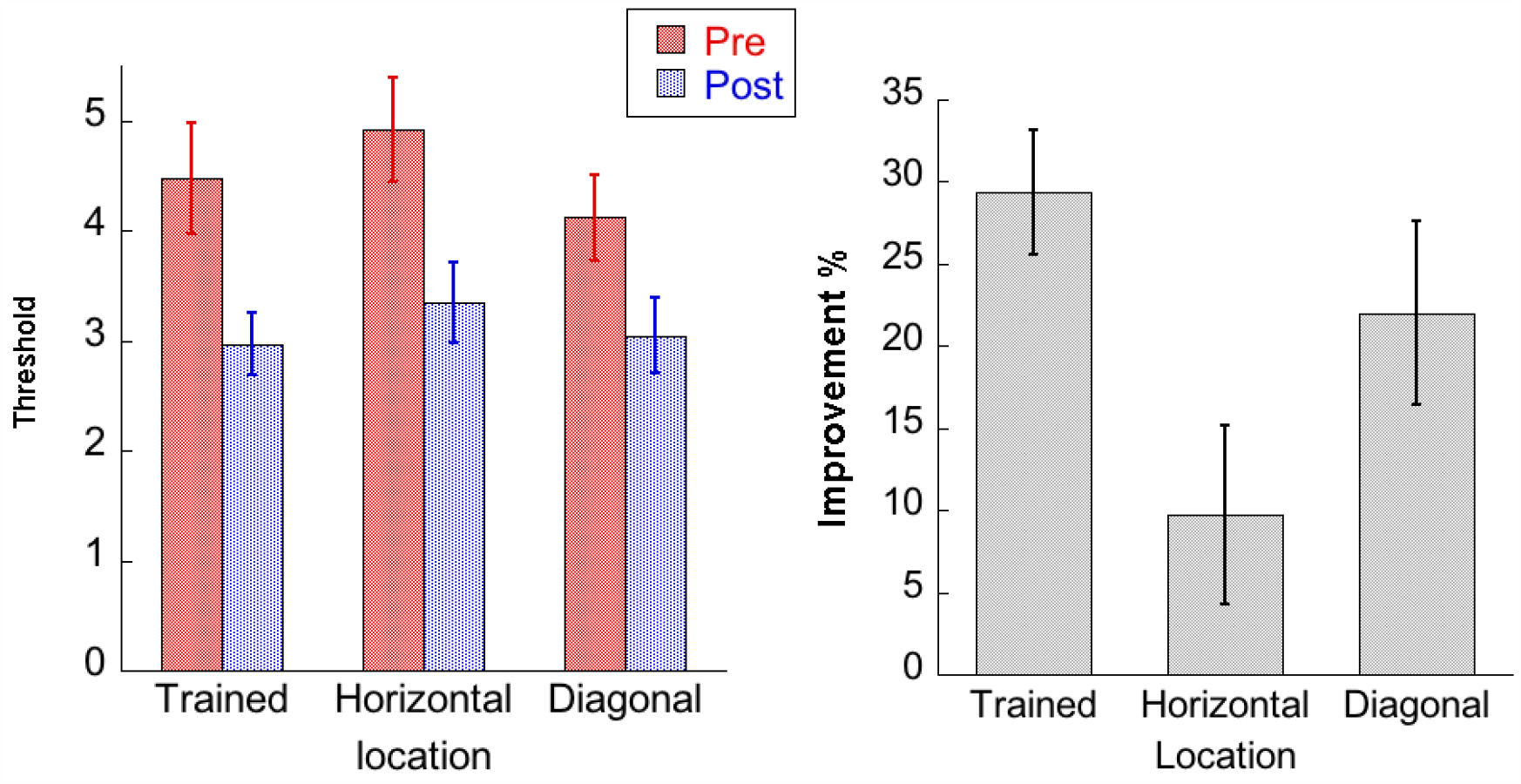
The result of Experiment 2

## Discussion

In the present study, we investigated the effect of a geometrical relationship between the trained location and an untrained location on transfer of VPL. In Experiment 1, we found that transfer occurs from the trained to a diagonal location, whereas no transfer was observed between the trained and a horizontal location. Since in Experiment 1, the diagonal line connecting the trained and diagonal locations passed the central fixation point, in Experiment 2, we examined whether the diagonal line needs to pass the fixation point. The result showed that transfer occurred along a diagonal line that did not pass the fixation point. In addition, it was found that transfer was obtained along a horizontal line passing the fixation point, although transfer did not occur along the horizontal line that did not pass the fixation point in Experiment 1. The results from Experiments 1 and 2 collectively suggest that transfer occurs either diagonally or along a line passing the fixation point.

To our knowledge, there is no evidence that feedback from higher-level stages provides biased enhancement along the diagonal line or lines passing the fixation point. On the other hand, activity in the extra striate cortex was enhanced when stimuli are symmetrically (Sasaki et al. 2005) or concentrically arranged around the fixation point (Wilson et al., 1997, Wilkinson et al., 2000). This suggests that our results may occur due to influences of mid-level visual stages.

In summary, we examined the effect of a geometric relationship between the trained location and an untrained location. We found transfer from the trained location along a diagonal line or a horizontal line passing the fixation point. These geometric-relationship specific transfers suggest that the specificity of VPL is at least partially determined by mid-level visual stages.

## Acknowledgements

This study was supported b**y** NSF-BSF BCS-2241417, NIH R01-EY031705, NIH R01-EY019466, NIH R01-EY027841.

## References

Andersen, G. J., Ni, R., Bower, J. D., & Watanabe, T. (2010). Perceptual learning, aging, and improved visual performance in early stages of visual processing. Journal of vision, 10(13), 4–4.

Ball, K., & Sekuler, R. (1987). Direction-specific improvement in motion discrimination. Vision research, 27(6), 953–965.

Bang, JW., Shibata, K. Frank, SM., Walsh, EG., Greenlee, MW., Watanabe, T., & Sasaki, Y. Consolidation and reconsolidation share behavioural and neurochemical mechanisms. Nature Human Behaviour 2 (7), 507–513

Byers, A., & Serences, J. T. (2014). Enhanced attentional gain as a mechanism for generalized perceptual learning in human visual cortex. Journal of neurophysiology, 112(5), 1217–1227.

Chang, L. H., Shibata, K., Andersen, G. J., Sasaki, Y., & Watanabe, T. Age-related declines of stability in visual perceptual learning. Current Biology 24 (24), 2926–2929

DeLoss, D.J., Watanabe, T., Andersen, G. J. (2015) Improving vision among older adults: behavioral training to improve sight. Psychological science 26 (4), 456–466

Dosher, B. A., & Lu, Z.-L. (1998). Perceptual learning reflects external noise filtering and internal noise reduction through channel reweighting. Proceedings of the National Academy of Sciences of the United States of America, 95(23), 13988–13993.

Fahle, M., & Poggio, T. (2002). Perceptual learning. MIT Press.

Frank, S. M., Becker, M ., Qi Geiger, P., Frank, U.I., Rosedahl, L.A., Malloni, W.M., Sasaki, Y.,Watanabe, T. (2022). Efficient learning in children with rapid GABA boosting during and after training. Current Biology 32 (23), 5022–5030. e7

Seitz, A. R., Nanez, J.E., Holloway, S.R., Watanabe, T. (2005). Visual experience can substantially alter critical flicker fusion thresholds. Human Psychopharmacology: Clinical and Experimental 20 (1), 55–60

Gilbert, C. D., Sigman, M., & Crist, R. E. (2001). The neural basis of perceptual learning. Neuron, 31(5), 681–697.

Gold, J. I., & Watanabe, T. (2010). Perceptual learning. Current Biology, 20(2), R46–R48.

Harris, H., Gliksberg, M., & Sagi, D. (2012). Generalized perceptual learning in the absence of sensory adaptation. Current Biology, 22(19), 1813–1817.

Hung, S. C., & Seitz, A. R. (2014). Prolonged training at threshold promotes robust retinotopic specificity in perceptual learning. The Journal of Neuroscience, 34(25), 8423–8431.

Karni, A., & Sagi, D. (1991). Where practice makes perfect in texture discrimination: evidence for primary visual cortex plasticity. Proceedings of the National Academy of Sciences, 88(11), 4966–4970.

Law, C. T., & Gold, J. I. (2008). Neural correlates of perceptual learning in a sensory-motor, but not a sensory, cortical area. Nature neuroscience, 11(4), 505–513.

Lemon, C., DeLoss, D., & Andersen, G. (2016). Improving collision detection in older adults using perceptual learning. Journal of Vision, 16(12), 541–541.

Levi, D. M., & Li, R. W. (2009). Perceptual learning as a potential treatment for amblyopia: a minireview. Vision research, 49(21), 2535–2549.

Levi, D. M., & Polat, U. (1996). Neural plasticity in adults with amblyopia. Proceedings of the National Academy of Sciences, 93(13), 6830–6834.

Liang, J., Zhou, Y., Fahle, M., & Liu, Z. (2015). Limited transfer of long-term motion perceptual learning with double training. Journal of vision, 15(10), 1–1.

Liang, J., Zhou, Y., Fahle, M., & Liu, Z. (2015). Specificity of motion discrimination learning even with double training and staircase. Journal of vision, 15(10), 3–3.

Mednick, S. C., Arman, A. C., & Boynton, G. M. (2005). The time course and specificity of perceptual deterioration. Proceedings of the National Academy of Sciences of the United States of America, 102(10), 3881–3885.

Roelfsema, P. R., van Ooyen, A ., Watanabe, T. (2010). Perceptual learning rules based on reinforcers and attention. Trends in cognitive sciences 14 (2), 64–71

Sagi, D., & Tanne, D. (1994). Perceptual learning: learning to see. Current opinion in neurobiology, 4(2), 195–199.

Sasaki, Y., Vanduffel, W., Knutsen, T., Tyler, C., & Tootell, R. (2005). Symmetry activates extrastriate visual cortex in human and nonhuman primates. Proceedings of the National Academy of Sciences of the United States of America, 102(8), 3159–3163.

Sasaki S, Watanabe T. Perceptual Learning. In: Werner J, Chalupa L, editors. The New Visual Neurosceinces. Cambridge: MIT Press; 2013. pp. 991–1000.

Sasaki, Y., Nanez, J., & Watanabe, T. (2010) Advances in visual perceptual learning and plasticity. Nature Reviews Neuroscience, 11(1):53–60.

Seitz, A. R., & Watanabe, T. (2003). Psychophysics: Is subliminal learning really passive?. Nature, 422(6927), 36–36.

Schoups, A. A., Vogels, R., & Orban, G. A. (1995). Human perceptual learning in identifying the oblique orientation: retinotopy, orientation specificity and monocularity. The Journal of physiology, 483(3), 797–810.

Schoups, A., Vogels, R., Qian, N., & Orban, G. (2001). Practising orientation identification improves orientation coding in V1 neurons. Nature, 412(6846), 549–553.

Shibata, K. W atanabe, T . Sasaki, Y. & Kawato, M. (2011). Perceptual learning incepted by decoded fMRI neurofeedback without stimulus presentation. Science 334 (6061), 1413–1415

K Shibata, LH Chang, D Kim, JE Náñez Sr, Y Kamitani Sasaki Y. & Watanabe, T. (2012). Decoding reveals plasticity in V3A as a result of motion perceptual learning. Public Library of Science 7 (8), e44003

Shibata, K., Sasaki, Y., Bang, J. W., Walsh, E. G., Machizawa, M. G., Tamaki, M., … & Watanabe, T. (2017). Overlearning hyperstabilizes a skill by rapidly making neurochemical processing inhibitory-dominant. Nature Neuroscience.

Tsushima, Y & Watanabe, T. (2009). Roles of attention in perceptual learning from perspectives of psychophysics and animal learning. Learning & behavior 37, 126–132

Watanabe, T., Náñez, J. E., Koyama, S., Mukai, I., Liederman, J., & Sasaki, Y. (2002). Greater plasticity in lower-level than higher-level visual motion processing in a passive perceptual learning task. Nature neuroscience, 5(10), 1003–1009.

Yamada, T., Watanabe, T., Sasaki, Y. (2023). Plasticity–stability dynamics during post-training processing of learning. Trends in Cognitive Sciences, S1364–6613 (23).

Watanabe, T., & Sasaki, Y. (2015). Perceptual learning: toward a comprehensive theory. Annual review of psychology, 66, 197.

Wilkinson, F., James, T. W., Wilson, H. R., Gati, J. S., Menon, R. S., & Goodale, M. A. (2000). An fMRI study of the selective activation of human extrastriate form vision areas by radial and concentric gratings. Current Biology, 10(22), 1455–1458.

Wilson, H. R., Wilkinson, F., & Asaad, W. (1997). Concentric orientation summation in human form vision. Vision research, 37(17), 2325–2330.

Xiao, L. Q., Zhang, J. Y., Wang, R., Klein, S. A., Levi, D. M., & Yu, C. (2008). Complete transfer of perceptual learning across retinal locations enabled by double training. Current biology: CB, 18(24), 1922.

Yang, T., & Maunsell, J. H. (2004). The effect of perceptual learning on neuronal responses in monkey visual area V4. The Journal of Neuroscience, 24(7), 1617–1626.

Yotsumoto, Y., Watanabe, T. (2008). Defining a Link between Perceptual Learning and Attention. PLOS biology, 6(8), 1623–1626.

Yotsumoto, Y., Watanabe, T., & Sasaki, Y. (2008). Different dynamics of performance and brain activation in the time course of perceptual learning. Neuron, 57(6), 827–833.

Yotsumoto, Y., Chang, L. H., Watanabe, T., & Sasaki, Y. (2009). Interference and feature specificity in visual perceptual learning. Vision research, 49(21), 2611–2623.

Yotsumoto, Y., Chang, L. H., Ni, R., Pierce, R., Andersen, G. J., Watanabe, T., & Sasaki, Y. (2014). White matter in the older brain is more plastic than in the younger brain. Nature communications, 5.

Zhang, T., Xiao, L. Q., Klein, S. A., Levi, D. M., & Yu, C. (2010). Decoupling location specificity from perceptual learning of orientation discrimination. Vision research, 50(4), 368–374.

